# An automated convergence diagnostic for phylogenetic MCMC analyses

**DOI:** 10.1101/2023.08.10.552869

**Authors:** Lars Berling, Remco Bouckaert, Alex Gavryushkin

## Abstract

Assessing convergence of Markov chain Monte Carlo (MCMC) based analyses is crucial but challenging, especially so in high dimensional and complex spaces such as the space of phylogenetic trees (treespace). In practice, it is assumed that the target distribution is the unique stationary distribution of the MCMC and convergence is achieved when samples appear to be stationary. Here we leverage recent advances in computational geometry of the treespace and introduce a method that combines classical statistical techniques and algorithms with geometric properties of the treespace to automatically evaluate and assess practical convergence of phylogenetic MCMC analyses. Our method monitors convergence across multiple MCMC chains and achieves high accuracy in detecting both practical convergence and convergence issues within treespace. Furthermore, our approach is developed to allow for real-time evaluation during the MCMC algorithm run, eliminating any of the chain post-processing steps that are currently required. Our tool therefore improves reliability and efficiency of MCMC based phylogenetic inference methods and makes analyses easier to reproduce and compare. We demonstrate the efficacy of our diagnostic via a well-calibrated simulation study and provide examples of its performance on real data sets. Although our method performs well in practice, a significant part of the underlying treespace probability theory is still missing, which creates an excellent opportunity for future mathematical research in this area.

The open source package for the phylogenetic inference framework BEAST2, called ASM, that implements these methods, making them accessible through a user-friendly GUI, is available from https://github.com/rbouckaert/asm/. The open source Python package, called tetres, that provides an interface for these methods enabling their applications beyond BEAST2 can be accessed at https://github.com/bioDS/tetres/.

## I. Introduction

Bayesian inference via the Markov chain Monte Carlo (MCMC) algorithm is widely used [1]–[4] to reconstruct the evolutionary relationships among a set of biological sequences. This inference paradigm utilizes probability distributions to characterize the uncertainty associated with all unknown parameters, including the evolutionary tree. Prior to observing the data, a probability distribution is assigned based on existing knowledge, known as the prior distribution. By applying Bayes’ theorem, one can calculate the probability of observing the sequence data given a specific tree topology and model parameters, resulting in the posterior distribution. MCMC algorithms are typically employed to approximate the posterior probability distribution, with various diagnostic techniques applied to assess the quality of this approximation.

The accuracy and precision of results from an analysis are dependent on the convergence of the MCMC samples to the target distribution, making it critical to ensure that this has been achieved [21]. We do not refer to theoretical convergence here, which is only guaranteed asymptotically as the number of samples approaches infinity. For the assumptions made in this manuscript, please refer to the end of the introduction. Convergence assessment remains a challenging and complex issue, not only in phylogenetics [5]–[13] but also the classical Euclidean statistics [14]–[20] In the context of phylogenetic MCMC, the state space is the space of all possible different evolutionary histories, a treespace, along with tree-associated Euclidean parameters, i.e. real-valued parameters that can be analysed using standard Euclidean statistical methods. Treespaces, being high-dimensional non-Euclidean spaces, pose the main and unique challenge in analysing phylogenetic data. The complexity of probability distributions over this already complex space further adds to the intricacy of this problem [22]–[27] Therefore, there is a need for practically effective methods to evaluate convergence of MCMC chains within the treespace to ensure the reliability of the resulting estimates and evolutionary conclusions.

Currently the most popular tool for phylogenetic MCMC convergence assessment is Tracer [8], which reads a log file containing a sample of trees and other parameters and produces effective sample size (ESS) estimates and trace plots for all Euclidean parameters associated with the analysis. However, convergence in this low dimensional Euclidean parameter space does not imply convergence in the high dimensional treespace or vice versa [17]. It is therefore crucial to have an assessment of convergence in both the treespace and the Euclidean parameter space associated with trees. Generally, classical methods to assess convergence assume a Euclidean parameter space and can therefore not be applied to phylogenetic trees directly.

To address these issues, multiple different approaches to represent and analyse samples of trees as a Euclidean parameter trace have been developed as an aid to assess the tree parameter convergence. The approach developed by Nylander, Wilgenbusch, Warren, *et al*. [5] uses split frequencies and branch lengths to adapt classical convergence assessment tools on samples of trees. The average standard deviation of split frequencies (ASDSF) [28] is another popular convergence assessment tool based on splits, i.e. the partition of taxa into two subsets. Furthermore, Lanfear, Hua, and Warren [6] and Warren, Geneva, and Lanfear [7] proposed a different approach to estimate the ESS of a sample of trees by utilising a vector of tree distances as a continuous parameter trace, referred to as the pseudo ESS. Yet another recently developed approach by Guimarães Fabreti and Höhna [11] converts trees into sequences based on the absence and presence of splits.

Despite these efforts, the problem of assessing convergence of the sample of trees directly remains largely unsolved and often overlooked in practice. A promising approach to solve this issue was discussed by Whidden and Matsen [26] in the form of a topological Gelman-Rubin diagnostic. The Gelman-Rubin diagnostic [16], [29], commonly known as the potential scale reduction factor (PSRF), is a widely used convergence assessment tool in MCMC analyses [18]. The diagnostic calculates a factor of the variances within and between multiple MCMC chains. This factor quantifies the potential improvement in our estimates with further sampling and indicates the degree of similarity among the independent samples. The *MrBayes* software [1] uses this diagnostic for continuous parameters, while the topological version developed in [26] cannot be used in practice due to prohibitive computational costs (NP-hardness) of the underlying distance measure between trees.

In this paper, we advance the idea of a tree Gelman-Rubin diagnostic (GR_*T*_), utilising the recently developed ranked nearest neighbour interchange (RNNI) treespace [30]–[34]. Our approach is structured in two parts: firstly, we develop the GR_*T*_ diagnostic for samples of trees. In the context of MCMC, samples of trees should exhibit near-indistinguishability between independent chains if drawn from the same distribution over the treespace. Our presented tree PSRF value quantifies this property of similarity. Building upon this foundation, we integrate our GR_*T*_ diagnostic into a comprehensive convergence assessment tool that incorporates continuous parameter trace estimates as well. This combination of components results in a powerful and robust convergence assessment diagnostic for phylogenetic MCMC analysis that advances the state of the art in the following ways.

Our approach is founded on a metric treespace that relies on tree rearrangement operations with efficiently computable distances [33], a unique feature not found in previous such treespaces. One advantage of these treespaces lies in the interpretability of the rearrangements they represent. In prior studies [32], [34], we demonstrated the potential of this particular treespace for statistical analysis of phylogenetic trees. Building upon this foundation, we utilize the concept of Fréchet variance, which allows a quantitative comparison of multiple distributions within the treespace. By utilizing a computationally efficient treespace for distance calculations we address previous limitations, namely the approach presented by Whidden and Matsen [26]. This paper introduces novel formulas for quantitatively assessing convergence and mixing within the treespace. Through rigorous evaluation in both simulations and real datasets, we demonstrate the necessity for such a tool to assess the convergence of tree samples. In addition, we automate the utilization of existing tools, ensuring fewer errors occur during the manual evaluation of these diagnostics in practice. Therefore, our approach quantifies the evaluation entirely, removing the necessity for visual or manual inspection and streamlining the process, thereby reducing the manual effort usually required for configuring multiple analyses. Additionally, we offer an easily accessible tool for practitioners in BEAST2, along with a Python implementation. We structure our paper as follows: first, we introduce our definition of a PSRF value for trees and demonstrate how it serves as the foundation for our GR_*T*_ diagnostic. In the Validation section, we provide evidence of the efficacy of our GR_*T*_ diagnostic by showing its performance on samples of trees from a well-calibrated simulation study. We then incorporate continuous parameter traces in our convergence assessment tool. In the Results section, we present the performance of our tree-based GR_*T*_ diagnostic on the widely used benchmark data sets DS1–DS11. Subsequently, we demonstrate the performance of the more sophisticated version on the same data sets, followed by an application to three real datasets. In conclusion, we highlight the promising implications of the RNNI treespace and the diagnostic for advancing statistical tools in the analysis of samples of phylogenetic time trees.

### Convergence

In theory, an ergodic MCMC is guaranteed to reach its unique stationary distribution when the number of samples/iterations reaches infinity. In practice this asymptotic property is never reached and we instead consider practical convergence. Loosely speaking we consider an MCMC analysis to be converged (as in practical convergence) if two (or multiple) independent MCMC analysis return indistinguishable collections of samples. More specifically, we assume that there exists a unique stationary distribution for our MCMC which is also the target distribution. Further, we also assume that if multiple independent MCMCs reach the same (or similar enough) distribution(s), i.e. essentially indistinguishable collections of trees, that they have (practically) converged to this stationary distribution.

Given our assumptions, we do acknowledge the possible challenge of pseudo-convergence and we revisit it in Section V. Discussion. However, detecting such a problem reliably in practice is generally impossible (unless the true distribution is known) so we do not explore this important topic in this paper. In this manuscript we will use the term convergence as an equivalent to practical convergence as described above.

## II. Methods

We present a tool that quantifies the similarity of multiple collection of trees, which in the context of MCMC samples can be used to assess whether independent MCMC runs are sampling the same tree distribution. We apply this novel tool to MCMC convergence assessment and present additional tools, based on existing practises in the field of phylogenetic MCMC, that aid in this endeavour. All quantities that we are estimating and comparing are sample quantities, including *V ar*_*T*_ (*t*). Pseudo-code for the individual algorithms presented is available in the Supplement Section S2 and implementations are available as a BEAST2 (Java) package [35] or as part of a Python package [36].

### A. Gelman-Rubin like diagnostic

For the remainder of the paper we will assume that the length of two (or multiple) chains is the same to simplify notations. Additionally, we presume that the variance for trees, a crucial component of our PSRF value, is finite. As demonstrated by Vehtari, Gelman, Simpson, *et al*. [20], the traditional 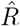 diagnostic fails when used in scenarios with infinite variance. It is imperative to note that in the realm of phylogenetic treespaces, our understanding of these quantities remains largely elusive.

The Gelman-Rubin diagnostic, sometimes also referred to as the PSRF or 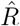 [29], is one of the most popular tools for diagnosing and monitoring convergence in MCMC algorithms [18]. The diagnostic is calculated as the ratio of the between-chain variance to the average within-chain variance. For the monitoring of convergence this ratio describes the factor by which we would expect the inferred samples to improve if the sampling process would be continued in the limit, i.e. the number of samples → ∞. With an increasing number of samples the PSRF value declines to 1.0 and a value much larger indicates that one or both of the following statements is true:

- Further sampling will substantially reduce the within chain variances, making the overall inference of the target distribution for each MCMC more precise
- Further sampling will substantially increase the between chain variances, indicating that the individual MCMCs have not completely sampled the target distribution

In either case we can conclude that our samples are far from convergence to the stationary distribution and we have to continue the sampling process. Hence, the PSRF can be seen as an indication of when the between-chain variance matches the average within-chain variance, implying sufficient similarity between the independently sampled distributions.

In this paper we introduce a diagnostic tool that we refer to as the GR_*T*_ diagnostic for evaluating convergence and termination of phylogenetic MCMC tree inference. This tool is inspired by the original diagnostic [29] as well as the topological version developed by Whidden and Matsen [26]. Due to the lack of a sample mean for phylogenetic trees we instead compute the within and between chain Fréchet variance using the RNNI tree metric [32], [33]. Specifically, we define the normalized Fréchet variance (a sample variance) of a tree *t* and a collection of trees *T* via the RNNI metric *d*(·,·) as

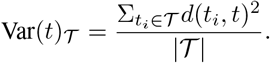

Since we assume all samples of trees to have the same size the normalization by the size of the set *T* is only relevant in practice when samples of trees have different sizes.

Using this variance we define our diagnostics PSRF value for a tree *t* given two sets of trees 𝒯 _1_ and 𝒯 _2_ as the square root of the ratio of variances for the tree *t* in the two sets.

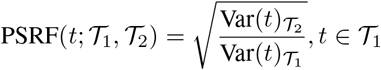

This definition differs from the method proposed in [26] by not using a 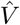 value that combines the between and in chain variances to a single value. Instead, our definition directly compares the tree’s Fréchet variance within its chain to its Fréchet variance in another chain, where the latter can be interpreted as the between chain variances akin to the original definition.

The overall PSRF value can be interpreted similarly to the originally proposed diagnostic with values close to 1.0 indicating indistinguishable tree sets and values much larger than 1.0 as different tree sets. In addition, our proposed PSRF value can also be less than 1.0 which is the case if the variance for a tree is lower in the set it does not originate from. This can be thought of as one tree set being “fully contained” within the perimeter of another tree set. Therefore, a convergence criterion in this case requires specifying a tolerance around the 1.0 value, which we will discuss below.

#### a) Generalisation for N independent chains

The diagnostic presented by Gelman and Rubin [29] was generalised by Brooks and Gelman [16] to incorporate more than 2 independent chains. We chose to initially present the construct using two chains for its simplicity of notation. Besides, using our slightly different definitions we can adapt these to the case of *N* independent chains as follows

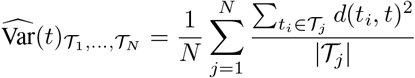

With this new notation of the between chain variance the PSRF value would be computed as

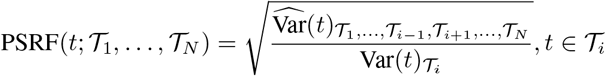

Note that increasing the number of independent chains will significantly increase computational cost as the pairwise distances between all combinations of chains and trees need to be calculated (and stored), i.e. for *N* independent chains of length *M* the number of necessary pairwise tree distance computations will be 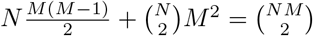.

#### b) Convergence assessment

Here we demonstrate the integration of tree PSRF values into our GR_*T*_ diagnostic. Unlike the original approach, we compute an average of PSRF values termed the GR value, considering the entire sample of trees from both chains. This is because we found that simply assessing the PSRF value is not sufficient (see Supplement Subsection S3-B) and it is important to put each iteration into the context of the full sample of trees from both (all) chains.

Given two MCMC chains and their respective samples of trees 𝒯_1_ and 𝒯_2_ up to the *i*-th sample, i.e. |𝒯_1_| = |𝒯_2_| = *i*, we define the GR value of tree 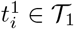 as

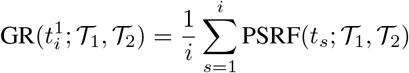

and analogously for tree 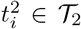. When calculating the GR value for sample *i* we have to newly calculate the PSRF value for each sample 1, …, *i* with the tree set containing all of the *i* samples and cannot reuse previously calculated PSRF values. However, this does not imply a significant increase in complexity because the required distances can be stored.

In the original diagnostic, it suffices for the PSRF value to fall below a threshold close to 1.0, typically set to 0.1 or lower, as suggested by Gelman, Carlin, Stern, *et al*. [37]. However, recent research by Vats and Knudson [19] challenges this threshold, indicating that 1.1 may be inadequate while also proposing new methods to enhance the robustness of the original diagnostic and determine an appropriate tolerance. Within this decision for chain termination lies the major difference of how we use the PSRF value. Through experimentation and practical application, we have found that a tolerance of 0.05 or less is suitable for our tree PSRF (see Supplement Subsection S3-A).

However, the main difference is not this stricter threshold but the requirement of the chain to consistently sample trees within this tolerance around 1.0 until a predetermined ESS estimate for the samples is reached. We utilize the RNNI distance and the median pseudo ESS variant for this estimation, which considers all trees in a sample as reference trees (see [10]). By requiring the chain to sample within the tolerance we ensure that the samples of trees from both (or all) chains are indistinguishably. If a GR value of a tree in either chain is not within the specified tolerance around 1.0 the samples up to this iteration get added to the tree burn-in and subsequently discarded from the set of trees that our assessment returns. Therefore, the diagnostic will return a subset of trees that has a sufficient pseudo ESS estimate and each tree in either chain has GR value within the specified tolerance around 1.0.

For a precise definition of the pseudo ESS, please refer to the Supplement Subsection S2-A and to [6], [10].

### B. The ASM package diagnostic

In addition to our described tree-based diagnostic, we introduce other diagnostics in our BEAST2 implementation [35]. These are aimed at enhancing the robustness of our diagnostic approach, particularly by incorporating continuous parameter traces. The first addition is an automated burn-in detection method, loosely inspired by Geweke [14], but to the best of our knowledge no such approach is used anywhere in practice. The second strategy that we incorporate is the concatenation of continuous parameter traces from the individual chains. We conclude the section with a high level description of the resulting comprehensive diagnostic.

#### a) Burn-in detection

As previously highlighted, our approach draws inspiration from the Geweke diagnostic [14], namely comparing samples from the start of a chain to the ones at the end of the chain to assess burn-in. However, to the best of our knowledge, our method is distinct from any existing practice. Given a continuous parameter trace we take its final quarter to calculate a mean *µ* and standard deviation *σ* and define a confidence interval as [*µ* − *σ, µ* + *σ*]. The algorithm iterates over the whole trace with a sliding window of width *M* = 10 until the window’s mean falls within this confidence interval. It then discards everything before this point as burn-in. In practice this diagnostic can be run for a set of specified parameters, which by default we set to be the posterior, likelihood, and prior. The overall burn-in is decided by the maximum value among all of these traces. The value *M* for averaging is aiding in making this detection more robust to random outliers as well as temporarily stuck chains.

#### b) Concatenation of traces

The second strategy we incorporate to automate our diagnostic involves using ESS thresholds for continuous parameter traces. Instead of applying these thresholds to the individual chains, we concatenate the traces before estimating the ESS. This concatenation offers several advantages. Firstly, it consolidates the MCMC output into a single collection of samples, thereby enhancing overall efficiency by doubling the total number of samples (in the case of two independent chains). Moreover, the concatenation process can potentially identify cases where individual chains sample parameters from different distributions. This is because concatenated traces from different distributions typically yield low ESS estimates. Indeed, it is important to note that while ESS was not designed for this purpose, this capability emerges as a by-product of the concatenation process. Similarly to the burn-in detection, we enforce this threshold only for the specified subset of priority parameters (posterior, likelihood, and prior by default).

So at the high level our algorithm can be described as seen in Algorithm 1.

##### Algorithm 1

High level pseudo code description of our diagnostic

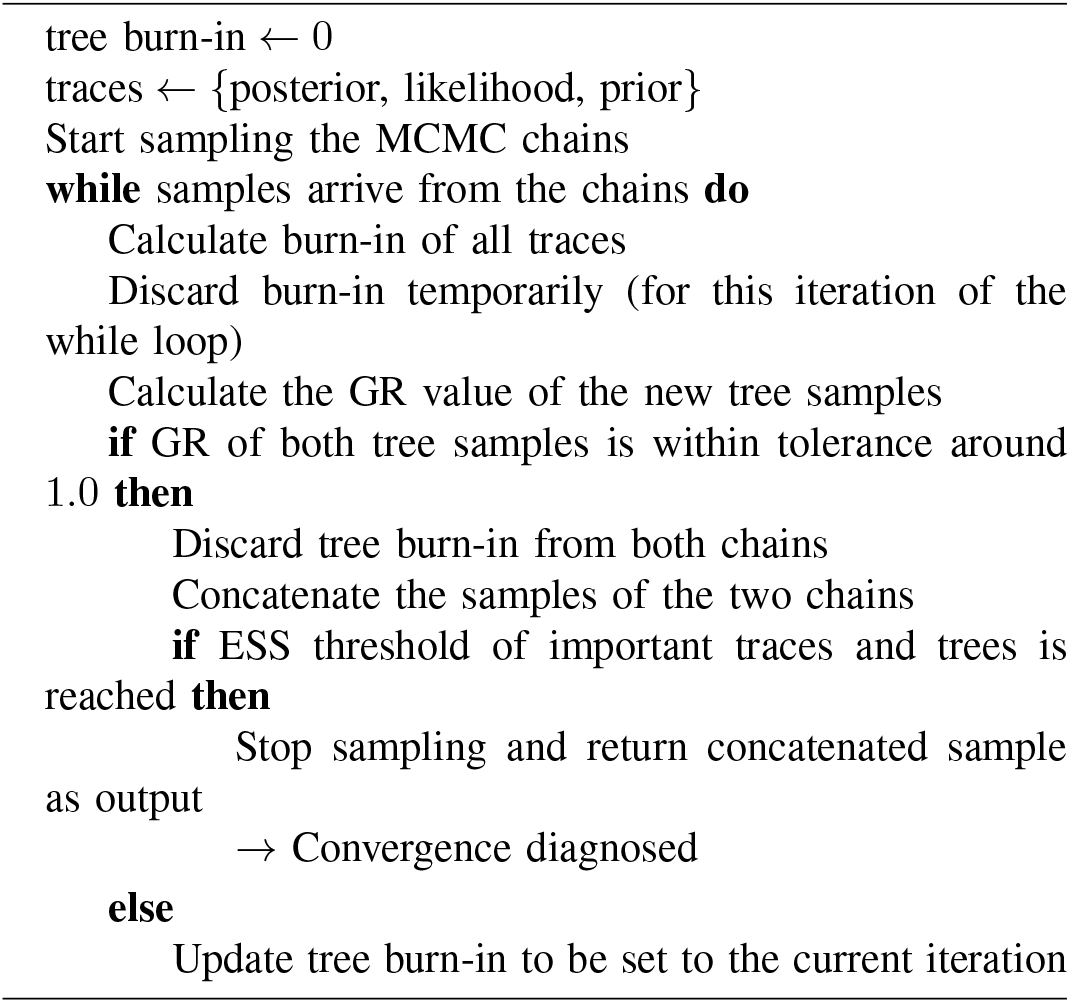

### C. Implementation

As previously mentioned, we have divided the content of this paper and our implementation into two distinct parts. The first part encompasses the purely tree based diagnostic, using the RNNI distance and is implemented into the time tree statistics python package *tetres* [36]. The main focus of this package is the implementation of statistical tools that are developed primarily over the RNNI treespace. In contrast, the second part is implemented within the BEAST2 framework as the ASM package [35] in the Java language.

## III. Validation

To validate our diagnostic, we conduct a well-calibrated simulation study. This section is again divided into two parts: firstly, we evaluate the treespace-based approach, and secondly, we repeat the evaluation for our comprehensive diagnostic implemented for BEAST2. For both diagnostics, we assess their performance on the well-calibrated simulation study, utilising regular ESS assessments, and conducting a coverage analysis.

### A. Well-calibrated simulation study

We performed a well-calibrated simulation study [38] sampling 50 tip dates randomly from the interval 0 to 1. We use a coalescent tree prior with constant population size lognormal(*µ* = −0.79, *σ* = 1.0) distributed giving a mean of 0.75, an HKY model with kappa log-normal(*µ* = 1, *σ* = 1.25) distributed, and gamma rate heterogeneity with four categories with shape parameter exponentially distributed with mean = 1 and frequencies Dirichlet(4, 4, 4, 4) distributed. Further, gamma is lower bounded by 0.1 [39]. We use a strict clock with uniform(0, 1) prior. Sampling 100 instances from this distribution using MCMC in BEAST2 [4], we get a range of tree heights from 1.03 to 14.74 with mean = 2.81 (note that due to the tips being sampled from 0 to 1, the tree height is lower bounded by 1) and we get a range for the clock rate of 0.01 to 0.99 in our study. With these trees, we sample sequences of 1000 sites using the sequence generator in BEAST2.

### B. Treespace based GR_T_ diagnostic

For the following results we used the implementation provided in [36] and MCMC analyses simulated as described in the previous section. This implies that we run our diagnostic on the full set of trees from an MCMC chain with sufficient burn-in discarded initially. For determining convergence, i.e. sufficiently many samples of trees with GR value close enough to 1.0, we used the pseudo ESS as described in the Supplement Subsection S2-A. The evidence provided in the first part of the section establishes the viability of our approach, serving as a proof of concept for our method.

#### a) Convergence

We compare different settings of our convergence diagnostic on the calibrated simulation study. One of the selected parameters that we compare are different pseudo ESS thresholds for the subset of trees, in this case 200 and 500. We choose these two values because 200 is a general rule of thumb in phylogenetic MCMC analyses [6] and 500 is because of recent findings by Magee, Karcher, Matsen IV, *et al*. [10] and Guimarães Fabreti and Höhna [11] which both advocate for a more stringent value higher than 200. In addition, we compare 3 different sizes of an acceptable interval (tolerance) for the GR value: 0.01, 0.02, and 0.05.

As we described in paragraph II-A0b we consider a pair of two independent chains converged (orange in Table I) if we find a large enough subset of trees in both chains that all have GR values within the respective tolerance around 1.0 and the pseudo ESS value of each subset of trees is at least the selected ESS threshold. If we reach the end of a chain without both of these conditions being met the two independent chains are considered not converged (0 and blue in Table I).

**TABLE I:** Our convergence diagnostic as implemented in tetres executed on 100 well-calibrated simulation studies with 1000 trees each.

| Tolerance | ESS threshold | Converged | Not converged |
| --- | --- | --- | --- |
| 0.05 | 200 | 100 | 0 |
|  | 500 | 100 | 0 |
| 0.02 | 200 | 99 | 1 |
|  | 500 | 92 | 8 |
| 0.01 | 200 | 80 | 20 |
|  | 500 | 53 | 47 |

We find that for the set of 1000 trees that arise from our simulation setup with a tolerance of 0.05 determines convergence in every case, whereas the more strict tolerances of 0.02 and 0.01 do not always detect convergence. The fundamental concept that underlies well-calibrated simulation studies is that an inference method employing the same model as the data’s generation should have the capability to accurately retrieve the true parameters. Therefore, these studies if setup appropriately are considered to return converged samples and are generally used to evaluate new methods and models [38]. Despite this notion, the fact that our diagnostic with strict tolerances does not detect convergence might imply that sampling only 1000 trees is sometimes insufficient even for such “nice” data. Therefore, in practice it might be useful to use a more strict tolerance to counter act premature convergence detection, particularly for more complex data sets.

Furthermore, it is important to highlight that the existing methods for assessing MCMC convergence [18] heavily rely on manual inspection guided by “expert” knowledge, which applies broadly to MCMC analyses in various fields, including phylogenetics. It is currently not possible to simulate scenarios that would demonstrate non-convergence without employing two chains on completely independent datasets. Therefore, we are constraint to use these well-calibrated simulation studies as our baseline to state that our diagnostic works as expected on datasets that would generally be considered converged analyses. This leaves a negative control as a non trivial open question for future research.

#### b) ESS assessment

As mentioned before, we did not use any continuous parameter trace ESS estimates to determine convergence but only used the pseudo ESS of the sample of trees using the RNNI distance. Here we check how the ESS estimates for all parameters behave when using our GR_*T*_ diagnostic. We find that in this well-calibrated setting the ESS estimates are in a similar range as the chosen ESS threshold value. These findings are displayed in Figure 1, where we compare the full MCMC chain versus the subset of tree samples determined by our GR_*T*_ diagnostic using a threshold of 200 or 500 respectively.

**Fig. 1:**
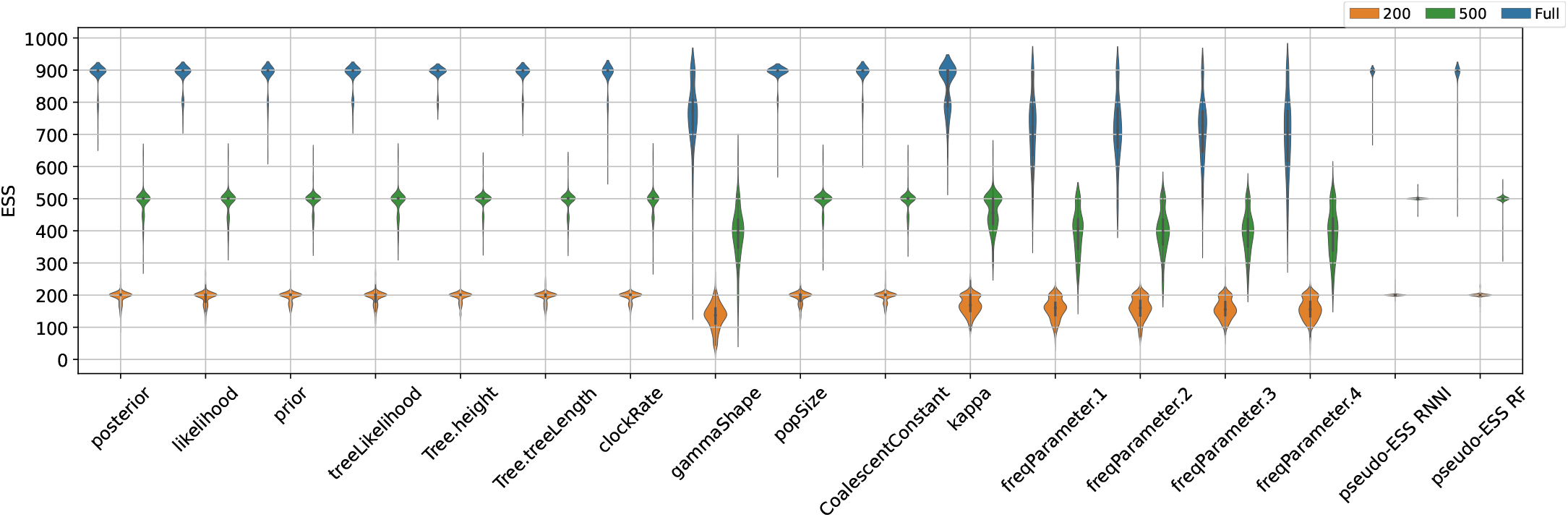
Comparing ESS estimates of the full chain versus the automatically determined portions using our diagnostic with RNNI pseudo ESS threshold values of 200 and 500 respectively. The figure additionally includes the RNNI and Robinson-Foulds (RF) pseudo-ESS estimates.

Moreover, we observe that in the case of our well-calibrated simulations the pseudo-ESS values, using RNNI or RF distances, align well with all other parameter ESS values after appropriately tuning the operator weights, see Figure 1. It remains open, whether the pseudo-ESS estimates can be used as a proxy for other ESS estimates or for calibrating MCMC operators in practice. Operators are used to propose new states for a corresponding parameter and the weight of an operator refers to its probability of being selected in each step of the MCMC. When a parameter *x* has low ESS at the end of an analysis it can be helpful to adjust the weights of operators to improve the sampling for this specific parameter. However, we emphasise that in a real scenario this strategy may not universally succeed due to mixing issues, the complexity of posteriors, or other reasons. This highlights one of the rationales behind our BEAST2 implementation’s utilisation of ESS estimates for the relevant continuous parameter, as elaborated in Subsection II-A.

#### c) Coverage analysis tetres

A coverage analysis for Bayesian MCMC refers to an evaluation of the accuracy and reliability of the posterior inference obtained trough the MCMC sampling methods. By comparing the true parameter values of a simulation with the parameters from the full posteriors (full MCMC chain) and the partial posteriors (subset of the full posterior) produced by our GR_*T*_ diagnostic, we are able to assess the accuracy and uncertainty of our method versus a manually set up MCMC run. The coverage is considered appropriate if it implies that the true parameter falls within the 95% confidence interval 95% of the time. This analysis helps to reveal any potential biases, under- or overestimation of uncertainties, allowing us to assess reliability of our diagnostic in comparison to a manually set up MCMC chain. For this coverage analysis we compare the full tree sets as sampled by the manually configured MCMC chain versus the subset of trees our diagnostic deems to be a convergent sample. The term “manually configured” denotes the best Bayesian phylogenetic practice of selecting the chain length, sampling intervals, and possibly burn-in before starting the MCMC analysis.

The results of this analysis are displayed in Table II. For a set of 100 independent simulations the 95% highest posterior density (HPD) of the binomial distribution ranges from 91 to 99 inclusive. As indicated by the red values (outside the aforementioned interval) our automatically determined sets of trees perform almost identical to the full simulation setups. It can be observed that using the ESS threshold value of 200 introduces low coverage in 3 additional parameters. However, we want to emphasise that low coverages are also affiliated with low ESS estimates, as visualised in Figure 1, implying insufficient sampling of the respective parameter. We touch upon this issue again in the next section with two ways to solve it in our BEAST2 implementation. Overall, this outcome gives us confidence that our GR_*T*_ diagnostic produces samples that are reliable estimates of the posterior and fully comparable to a manually set up MCMC analysis. As before the results are achieved with a GR tolerance of 0.05. More coverage analyses for different settings, including the use of the median instead of the mean for smoothing of the GR value, can be found in the Supplement, see Subsection S4-A.

**TABLE II:** Coverage of the true value by 95% HPD estimates from 100 independent simulations. Columns correspond to the pseudo ESS threshold of 200, 500, and the full sample of trees. The red coloured values indicate coverage outside the expected range between 91 and 99, the two values separated by ‘/’ represent the two independent chains.

| Parameter | Expected range (91-99) |  |  |
| --- | --- | --- | --- |
|  | 200 | 500 | Full |
| Tree.height | 93/93 | 96/96 | 96/96 |
| Tree.treeLength | 94/92 | 96/95 | 97/94 |
| clockRate | 91/91 | 90/92 | 91/93 |
| gammaShape | 87/90 | 93/93 | 92/92 |
| popSize | 90/93 | 97/96 | 96/97 |
| CoalescentConstant | 91/92 | 93/92 | 92/92 |
| kappa | 93/94 | 93/93 | 94/94 |
| freqParameter.1 | 95/92 | 92/92 | 92/92 |
| freqParameter.2 | 90/93 | 94/94 | 94/95 |
| freqParameter.3 | 86/87 | 89/87 | 87/90 |
| freqParameter.4 | 94/94 | 94/93 | 95/95 |

### C. BEAST2 ASM package validation

In the previous section we used a manually set up simulation that had fully run all the predetermined iterations. For the validation of our automated stopping implementation we reran these analyses using our automated convergence diagnostic package ASM and performed a coverage analysis for the resulting samples of trees.

#### a) Coverage analysis ASM

As displayed in Table III the coverage analysis is very similar to the previous one (Table II), slight difference are expected as MCMC is a random process. This result again confirms that our diagnostic produces reliable estimates of the posterior distribution.

**TABLE III:** Coverage of parameters when executing 100 well-calibrated simulations using the ASM package.

|  | coverage | mean ESS | min ESS |
| --- | --- | --- | --- |
| Tree.height | 95 | 224.70 | 57.56 |
| Tree.treeLength | 94 | 219.30 | 95.49 |
| clockRate | 91 | 222.71 | 66.45 |
| gammaShape | 89 | 124.27 | 16.35 |
| popSize | 91 | 222.54 | 95.05 |
| kappa | 92 | 178.38 | 86.39 |
| freqParameter.1 | 91 | 181.88 | 51.34 |
| freqParameter.2 | 94 | 178.08 | 52.26 |
| freqParameter.3 | 88 | 176.19 | 44.24 |
| freqParameter.4 | 96 | 182.90 | 41.98 |

**TABLE IV:** Different MCMC configurations used for DS1– DS11.

| Config | Samples | thinning | pre-burn-in |
| --- | --- | --- | --- |
| 1 | $2.5 \times 10^6$ | $2.5 \times 10^3$ | 0 |
| 2 | $2.5 \times 10^6$ | $2.5 \times 10^3$ | $1 \times 10^6$ |
| 3 | $5 \times 10^6$ | $5 \times 10^3$ | $1 \times 10^6$ |
| 4 | $5 \times 10^6$ | $5 \times 10^3$ | $2 \times 10^6$ |
| 5 | $1 \times 10^7$ | $1 \times 10^4$ | 0 |
| 6 | $1 \times 10^7$ | $5 \times 10^3$ | 0 |

A minor difference is the coverage of the frequency parameters, which do not quite reach the required 91-99 expected range. Additionally, the table also displays the minimum ESS estimate of these parameters across the 100 simulations, see column min ESS in Table III. For parameters with very low minimum ESS, such as the frequency parameters, a low coverage is expected because of insufficient number of samples. We want to emphasise that failing the coverage test is not a major concern as there are two ways to circumvent such a problem for any parameter in general using our tool. The first approach is that we can add any parameter in question to the list of important parameters in our diagnostic, which in turn would enforce a sufficiently high ESS estimate and increase the coverage. The second approach is to tune the operator weights in the analysis setup, which will also result in increased ESS estimates.

Furthermore, the inferred parameters of the sample are all highly correlated to the true values, this is visualised for the failed frequency parameter in Figure 2. This visualisation further indicates that there is no major concern about the low coverage parameters.

**Fig. 2:**
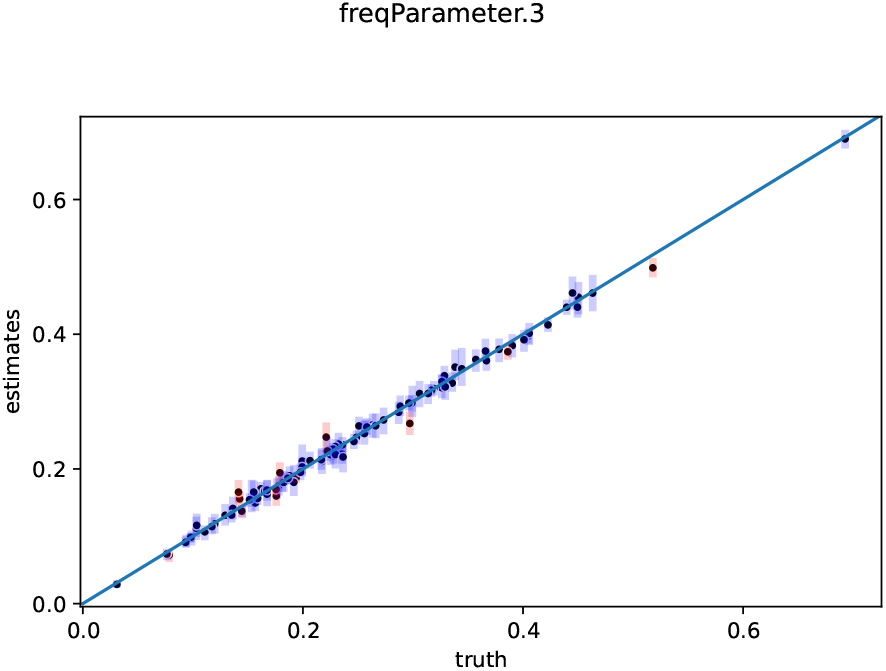
Visualising the coverage test for frequency parameter 3, the *x* axis displays the true parameter value and the *y* axis the inferred parameter. Blue boxes indicate that the true value is within the 95%HPD interval.

## IV. Results

We apply our convergence diagnostic technique to the datasets DS1–DS11, which are popular for benchmarking in computational phylogenetics [24]–[26], [28]. Previous findings on these suggest multi-modality in the treespace which implies that MCMC exploration and convergence within the treespace is challenging [24], [26]. We start by presenting the results of our tree based diagnostic first and find that they are coherent with these previously identified problems. Then, we present the results of applying our BEAST2 ASM package to the same datasets and find that the combination of continuous trace ESS estimates and our GR_*T*_ diagnostic performs well on these datasets. Additionally, we demonstrate the effectiveness of our package on three real datasets, revealing no major difficulties in identifying converged MCMC runs.

### A) Convergence diagnostic on DS1–DS11

For each of these 11 dataset, we set up 6 different BEAST2 analyses to generate samples of fully resolved time trees. Subsequently, our convergence diagnostic as implemented in tetres was applied to these 66 sets of trees. Additionally, we present results based on our BEAST2 package ASM, yielding comparable results, which among other things serves as a validation of the two implementations correctness. Based on these findings, we investigated the number of tree samples required to meet the specified pseudo ESS threshold (in this case 200). This investigation provides further insights into the mixing quality of the tree samples, as measured by pseudo ESS.

#### a) Configurations

We used the following 6 analysis configurations to generate two independent chains for each of the DS1–DS11 datasets.

In Table V we visualise the results of our diagnostic on these data sets. Indicated in cyan are runs that, according to our diagnostic, have not reached convergence yet, and could therefore indicate convergence problems within treespace. The table also contains the results of three different settings for the tolerance of our diagnostic. The diagnostic performs as expected with lower tolerances being less likely to reach convergence as it is stricter.

**TABLE V:** Assessing our diagnostic on DS1–DS11 with different MCMC setups. A cell labelled 1 (orange) indicates a run diagnosed as converged, a 0 (cyan) indicates that no convergence is diagnosed within the range of the sample. See Table IV for the different MCMC configurations C1,, C6.

|  | ESS-threshold | mean - 0.05 |  |  |  |  |  | mean - 0.02 |  |  |  |  |  | mean - 0.01 |  |  |  |  |  |
| --- | --- | --- | --- | --- | --- | --- | --- | --- | --- | --- | --- | --- | --- | --- | --- | --- | --- | --- | --- |
|  |  | C1 | C2 | C3 | C4 | C5 | C6 | C1 | C2 | C3 | C4 | C5 | C6 | C1 | C2 | C3 | C4 | C5 | C6 |
| DS1 | 200 | 0 | 0 | 1 | 1 | 1 | 1 | 0 | 0 | 0 | 1 | 1 | 1 | 0 | 0 | 0 | 0 | 1 | 1 |
|  | 500 | 0 | 0 | 0 | 0 | 0 | 1 | 0 | 0 | 0 | 0 | 0 | 1 | 0 | 0 | 0 | 0 | 0 | 0 |
| DS2 | 200 | 1 | 1 | 1 | 1 | 1 | 1 | 1 | 1 | 1 | 1 | 1 | 1 | 1 | 0 | 1 | 1 | 1 | 1 |
|  | 500 | 1 | 1 | 1 | 1 | 1 | 1 | 1 | 1 | 1 | 1 | 1 | 1 | 0 | 0 | 0 | 1 | 0 | 1 |
| DS3 | 200 | 1 | 1 | 1 | 1 | 1 | 1 | 1 | 1 | 1 | 1 | 1 | 1 | 1 | 1 | 1 | 1 | 1 | 1 |
|  | 500 | 1 | 1 | 1 | 1 | 1 | 1 | 1 | 1 | 1 | 1 | 1 | 1 | 1 | 1 | 1 | 1 | 1 | 1 |
| DS4 | 200 | 0 | 1 | 1 | 1 | 1 | 1 | 0 | 0 | 0 | 1 | 1 | 1 | 0 | 0 | 0 | 0 | 0 | 0 |
|  | 500 | 0 | 0 | 0 | 0 | 1 | 1 | 0 | 0 | 0 | 0 | 0 | 1 | 0 | 0 | 0 | 0 | 0 | 0 |
| DS5 | 200 | 1 | 1 | 1 | 1 | 1 | 1 | 1 | 1 | 1 | 1 | 1 | 1 | 0 | 1 | 1 | 1 | 1 | 1 |
|  | 500 | 1 | 0 | 1 | 1 | 1 | 1 | 0 | 0 | 1 | 1 | 1 | 1 | 0 | 0 | 1 | 1 | 0 | 1 |
| DS6 | 200 | 1 | 1 | 0 | 1 | 1 | 1 | 0 | 0 | 0 | 0 | 1 | 1 | 0 | 0 | 0 | 0 | 0 | 1 |
|  | 500 | 0 | 0 | 0 | 0 | 1 | 1 | 0 | 0 | 0 | 0 | 1 | 0 | 0 | 0 | 0 | 0 | 0 | 0 |
| DS7 | 200 | 1 | 1 | 1 | 1 | 1 | 1 | 1 | 1 | 1 | 1 | 1 | 1 | 1 | 1 | 1 | 1 | 1 | 1 |
|  | 500 | 1 | 1 | 1 | 1 | 1 | 1 | 0 | 0 | 1 | 1 | 1 | 1 | 0 | 0 | 0 | 0 | 0 | 1 |
| DS8 | 200 | 1 | 1 | 1 | 1 | 1 | 1 | 1 | 1 | 1 | 1 | 1 | 1 | 1 | 0 | 1 | 1 | 1 | 1 |
|  | 500 | 1 | 1 | 1 | 1 | 1 | 1 | 1 | 1 | 1 | 1 | 1 | 1 | 1 | 0 | 1 | 1 | 1 | 1 |
| DS9 | 200 | 0 | 1 | 1 | 1 | 1 | 1 | 1 | 1 | 1 | 1 | 1 | 1 | 1 | 1 | 1 | 1 | 1 | 1 |
|  | 500 | 0 | 1 | 1 | 1 | 1 | 1 | 0 | 0 | 1 | 1 | 1 | 1 | 0 | 0 | 1 | 1 | 1 | 1 |
| DS10 | 200 | 1 | 1 | 1 | 1 | 1 | 1 | 0 | 0 | 1 | 1 | 1 | 1 | 0 | 0 | 1 | 1 | 1 | 1 |
|  | 500 | 1 | 0 | 0 | 0 | 0 | 0 | 0 | 0 | 0 | 0 | 0 | 0 | 0 | 0 | 0 | 0 | 0 | 0 |
| DS11 | 200 | 1 | 1 | 1 | 1 | 1 | 1 | 1 | 1 | 1 | 1 | 1 | 1 | 1 | 1 | 1 | 1 | 1 | 1 |
|  | 500 | 0 | 1 | 1 | 1 | 1 | 1 | 1 | 1 | 1 | 1 | 1 | 1 | 1 | 1 | 1 | 1 | 1 | 1 |

Whidden and Matsen [26] found that DS1, DS4, DS5, DS6, DS7, and DS10 are peaky, meaning they contain multiple regions in treespace with high posterior probability which are separated by regions of low posterior probability and therefore harder to sample. In Table V it can be seen that our diagnostic agrees with these findings indicating that DS1, DS4, DS5, DS6, DS7 and DS10 are often unable to reach convergence in our setup. This indicates that these data sets have tricky to sample posterior distribution over treespace.

Our findings also show that DS2, DS3, DS8, DS9, and DS11 are reaching convergence in most of the setup runs. This indicates that the posterior distribution over treespace is easier to sample and hence reaches convergence with fewer samples. Again, this results is consistent with findings of Whidden and Matsen [26], who indicate that the distributions of DS2, DS3, and DS8 do not contain any peaks.

#### b) BEAST2 ASM package on DS1–DS11

We tested the BEAST2 implementation on the DS1–DS11 datasets ten times, utilising a strict clock with a fixed rate of 1, HKY substitution model, and Yule tree prior. The stopping criteria employed consist of GR_*T*_ statistic with a tolerance of 0.05, requiring pseudo ESS to exceed 200, and the ESS estimate of posterior, likelihood, and prior must also exceed 200. Table VI presents the average chain length, with the log frequency set to 10 thousand. As a result, the average trace and tree log for DS1, for example, contains 246.1 samples.

**TABLE VI:** Average, shortest, and longest chain lengths in millions of samples and fraction of longest divided by shortest chain length over 10 runs for data sets DS1–DS11. Lengths of the 10 individual runs are available in the Supplement Section S5.

| Data set | Average | Shortest | Longest | Longest/shortest |
| --- | --- | --- | --- | --- |
| DS1 | 2.461 | 1.99 | 3.73 | 1.87 |
| DS2 | 2.378 | 1.99 | 3.28 | 1.65 |
| DS3 | 3.435 | 2.32 | 4.36 | 1.88 |
| DS4 | 4.235 | 2.00 | 9.83 | 4.92 |
| DS5 | 2.454 | 2.00 | 3.41 | 1.71 |
| DS6 | 2.902 | 2.00 | 3.51 | 1.76 |
| DS7 | 3.797 | 2.34 | 4.67 | 2.00 |
| DS8 | 2.601 | 1.99 | 3.53 | 1.77 |
| DS9 | 2.697 | 1.99 | 4.35 | 2.19 |
| DS10 | 2.551 | 2.21 | 3.34 | 1.51 |
| DS11 | 2.579 | 2.00 | 3.57 | 1.79 |

**TABLE VII:** Average, shortest, and longest MCMC chain lengths over 10 runs before automatically stopping. Lengths of all 10 runs can be found in the Supplement Table S7.

|  | HCV | Primates | COVID-19 |
| --- | --- | --- | --- |
| average | 1474500 | 4340000 | 77364000 |
| shortest | 4650000 | 2030000 | 45630000 |
| longest | 19490000 | 11810000 | 148170000 |
| ratio | 4.19 | 5.82 | 3.25 |

We conducted visual inspection of the traces in Tracer [8] for all datasets, confirming convergence to the same posterior distribution. In all cases, there was a maximum difference of 2 between the lowest and highest log posterior, as well as prior and likelihood estimates across the ten runs, indicating that each run effectively converged to the same distribution. The inclusion of an ESS requirement of at least 200 for traces and trees leads to a minimum chain length of 200 samples (1.99 million steps), which, with a log frequency of 10 thousand, results in a minimum chain length of 2 million. The majority of chain lengths in the experiment were only slightly longer than 2 million, except for DS4, which exhibited two outliers just under 10 million (see Supplement Section S5). This variation can be attributed to the inherent stochastic nature of MCMC, causing differences in convergence times.

The presence of chains with “only” 200 samples suggest that the trace ESS criterion limits the stopping process, while the other criteria suggest that the MCMC could be terminated earlier. On the other hand, longer stopping times for some chains were accompanied by much higher ESS estimates for posterior, prior and likelihood, with the GR_*T*_ criterion being the primary factor in prolonging the sampling process. These findings indicate that neither criterion can be eliminated, as both are essential in determining the stopping point under different circumstances or conversely form a bottleneck in certain situations.

#### c) Further investigating mixing of trees

Due to the nature of MCMC analyses the resulting samples are not independent, hence the idea of an ESS measures to estimate the number of independent samples. Here we are further investigating this non-independence in treespace using our different configurations on the 11 data sets via the median pseudo ESS measure [6], [10]. Specifically, we are investigating how many samples are needed of a converged part of a chain to reach a desired pseudo ESS threshold,

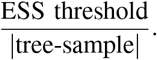

This value is closer to 1 if the MCMC thinning is appropriate, meaning that the samples are basically independent. However, smaller values indicate that the thinning interval is too small and therefore a larger number of samples is needed to reach the desired number of independent samples, i.e. the ESS threshold.

We visualised these findings in Figure 3 and it can be observed that the data sets DS1, DS4, and DS10 have more non-independent samples than others. In contrast, almost all samples in data sets DS2, DS3, and DS8 are independent (fraction close to 1). We also visualised these values for the samples resulting from our BEAST2 implementation in the ASM labelled column, which has very similar values among all datasets except DS3, DS4, and DS7.

**Fig. 3:**
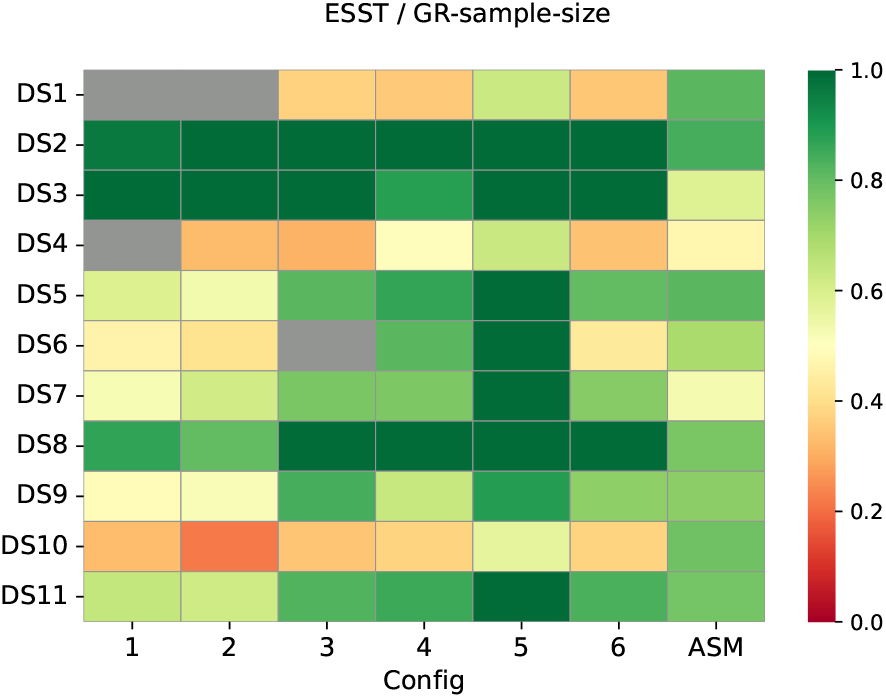
Fraction of ESS threshold over GR_*T*_ determined sample size. The grey cells indicate runs that are not considered converged. Dark green indicates that the sample size is the same as the ESS threshold (200 in this case). See Table IV for the different MCMC configurations on the x-axis. Accessible version available in the Supplement Figure S1.

We conclude from this investigation that the previously identified “peaky” datasets DS1, DS4, and DS10 display tree mixing that is coherent with these findings. However, we do not find stark differences among any of the datasets using our ASM package. Moreover, we do not find any scenarios that would suggest independent chains would be stuck in different modes. Rather, the datasets that are not diagnosed as converged have insufficient chain setups that sample too few iterations and trees. If this is indeed the case, it results in a simple fix of running the chain longer rather than having to investigate and deal with multimodal tree distributions.

### B. Real data application of convergence assessment

We first analysed a dataset comprising of 63 hepatitis-C virus (HCV) sequences sampled in Egypt in 1993, previously analysed in [40] and [41]. To avoid committing to a specific site model, we used model averaging with bModelTest [42], a strict clock, and Yule skyline [43] tree prior providing a flexible multi epoch pure birth model.

Furthermore, we analysed 36 mammalian species [44] again with bModelTest site model, an optimised relaxed clock [45], and a Yule tree prior together with a number of node calibrations used in [44].

Lastly, we examined an alignment of 257 full COVID-19 genome sequences, a subset from [46], using four partitions as in the original study with a HKY model each, estimated frequencies and relative substitution rate for each partition. Further, we used a strict clock and the BICEPS [42] tree prior, which provides a flexible multi epoch coalescent model. Unlike the previous datasets, this analysis incorporated dated tips.

Each analysis was run ten times with stopping criterion the same as in paragraph IV-A0b, and a log frequency of 10 thousand samples. For each data set, we verified convergence to the same parameter distribution across all chains through visual inspection using Tracer.

In the case of the HCV data, the shortest run took 4.65 millions steps and the longest extended just beyond 19.49 million steps, resulting in a ratio of 4.19 between the longest and shortest run. Similarly, the primate data exhibited a shortest chain length of 2.03 million steps and a longest of 11.81 million, yielding a ratio of 5.82. Chains on the COVID-19 data spanned from 45.6 to 148 million steps, returning a slightly smaller ratio of 3.25.

The notable disparity in convergence rates between fast and slow chains in real data may suggest an approach of starting multiple chains and selecting the fastest to reach convergence in order to accelerate analyses. However, it remains uncertain whether slow converging instances inefficiently sample the state space or merely become trapped in unexplored modes avoided by faster converging chains. Further investigation is required to establish whether this strategy will be suitable.

## V. Discussion

This paper presents a diagnostic tool for the automated assessment of convergence in phylogenetic MCMC analyses, utilising the RNNI treespace and its associated metric. Through an extensive evaluation on well-calibrated simulation studies, we demonstrate the efficacy of our diagnostic. We split the evaluation of our method into two separate parts. First, we demonstrate that our diagnostic tool is capable of producing reliable estimates of the posterior when applied only to samples of trees, without using any continuous parameters. Then, we extend this tool to incorporate continuous parameter traces. This comprehensive approach enables automatic convergence assessment of all parameters of interest in typical phylogenetic applications. We demonstrate this by applying our method to multiple real data sets.

This approach is a further advance in the area of developing statistical methods over geometrically complex spaces, such as the treespace, which has direct practical implications for phylogenetic MCMC analyses. By eliminating manual visualisation-based steps in setting up MCMC chains, our tool makes these analyses more accessible, reduces variability in the setup process, and contributes improving reproducibility of results obtained using phylogenetic MCMC analysis. In addition, automatic convergence diagnostic is a major stepping stone to enable online phylogenetic inference algorithms [47]. Yet to be explored applications of our tool include comparing tree samples inferred using different evolutionary models and advancing the “convergence” assessment and setup process when using nested sampling approaches [48].

While evaluating and developing this diagnostic we made a few peculiar observations that we believe are worth reporting. First, as we demonstrate on our well-calibrated simulation study, a sample of 1000 trees may not consistently guarantee convergence, particularly when tolerance is low. Second, as we analysed DS1–DS11 we found that the convergence assessment was coherent with previous findings on these data sets (specifically multimodality issues implying hard to sample posteriors). However we did not find cases of independent chains being stuck in different modes. Moreover, we have reason to believe that the unconverged runs can be attributed to chains being set up with inadequate lengths and sampling intervals. As demonstrated by runs using the ASM package on these DS1–DS11 datasets we do determine convergence for all runs eventually. Therefore, we conjecture that in these experiments a prolonging of the chain’s runtime would lead to convergence being diagnosed by our diagnostic.

Another aspect important in practice is the use of ESS estimates, specifically in cases when the log frequency is high and every sample is independent of the previous one. Note that when using phylogenetic MCMC, it is common practice to have high thinning values, i.e. it is common to log “only” 1000 samples from a chain that ran for 10 million iterations. However, in situations where the effective sample size and actual sample size will be very close to each other the ESS estimates are unstable. We observed cases where the ESS would double by adding a single sample, hence ESS based criteria work better when the number of independent samples is smaller than the number of MCMC samples. Hence, we infer that enhancing the robustness of the ESS estimators is necessary, although this is beyond the scope of this paper.

We finish this paper by outlining possible future directions as we see them and highlighting open problems that are yet to be addressed in the context of phylogenetic MCMC analyses and convergence assessment. Like we mentioned before, the sample of trees is the most important output of a phylogenetic MCMC analysis and should therefore be considered as such, especially when assessing convergence. However, it is an unsolved problem whether convergence in the treespace also implies convergence in other low dimensional parameter spaces. Our results in the well-calibrated simulation setting hint to this conjecture being true, but it would require major effort to show that this is indeed the case for any real scenario. Therefore, in practice it is still important to use more than one check for convergence which is why our implementation incorporates the use of continuous parameter traces in addition to the treespace based approach.

Another direction of ongoing research into probability distributions over treespaces is related to tree islands or terraces [22]–[24] [27]. This can also be thought of as multi-modal distributions, where high posterior probability regions in the treespace are separated by valleys of low posterior probability. It has previously been shown in [24], [26] that some of the data sets DS1–DS11 suffer from this phenomenon of multi modality, which can lead to false conclusions about an analysis. Our diagnostic is able to detect such problems in these case where independent chains are stuck in different parts of the treespace (see Supplement Subsection S3-A for more detail).

A problem that is related to multi-modality is pseudo-convergence, which is the phenomenon where an MCMC chain is apparently sampling from the stationary distribution but in reality it is stuck somewhere that is not in fact the stationary distribution. This can be the case when it takes much longer than the length of the chain to transition between this pseudo stationary part and the true stationary distribution. For example, when this may happen when the stationary distribution is multi-modal. However, multi-modality does not imply pseudo-convergence as sometimes the chain is able to transition between modes, or cross the valleys, and sample from the true stationary multimodal distribution. In fact, in phylogenetics multi-modality is a poorly explored phenomenon, due to geometric complexity of the treespace, with even the notion of a mode being somewhat esoteric. Altogether, the phenomena of multi-modality and pseudo-convergence form a compounded challenge.

These problems can potentially cause our diagnostic tools to converge to inaccurate conclusions. For example, if all independent chains pseudo-converged to the same mode, it is fundamentally impossible for a distance based method to diagnose pseudo-convergence. This and the problem of identifying multi-modal tree distributions remain open problems that need to be carefully addressed to deepen our understanding of distributions over the treespace. In this context, the approach developed in this paper provides additional insight into this largely unexplored area of phylogenetic MCMC.

## Data and software availability

Data used in the experiments can be accessed through https://github.com/rbouckaert/asm/releases/tag/v0.0.1. ASM is a BEAST 2 package that provides a user friendly GUI to the methods introduced here. It is open source and available at https://github.com/rbouckaert/asm. The Python package tetres provides an interface to the RNNI based methods introduced here. It is open source and available at https://github.com/bioDS/tetres.

## Acknowledgement

This work was partially supported by Royal Society Te Apārangi through a Rutherford Discovery Fellowship (UOC1702) and a Marsden grant (21-UOC-057), and by Ministry of Business, Innovation, and Employment of New Zealand through an Endeavour Smart Ideas grant (UOOX1912) and a Data Science Programmes grant (UOAX1932).

We thank Luiz Carvalho and the members of the bioDS lab for their feedback and discussion of the results.

## S. Accessible data visualisation

An accessible version of tree mixing plot Figure 3.

**Fig. S1:**
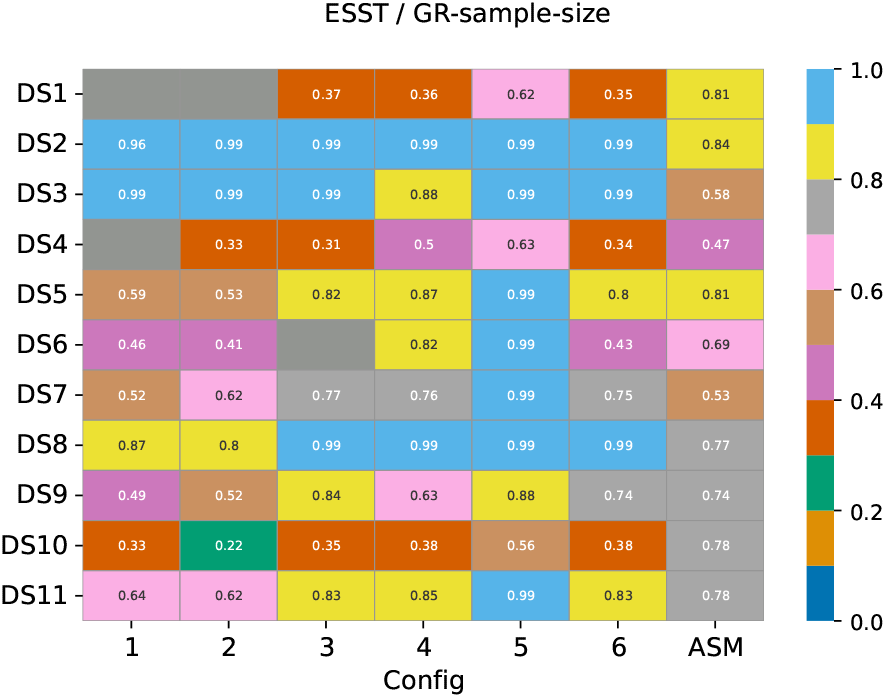
Accessible version of Figure 3. Labels in each rectangle display the value of the fraction.

## S. Pseudo code of presented algorithms

We begin by introducing all the preliminary algorithms and conclude with a comprehensive pseudo code of our diagnostic, offering more algorithmic depth than the version presented in the main paper.

### A. An effective sample size measure for a set of trees

In this manuscript we use the following definition of pseudo ESS [6], [10], which is the computation of an ESS measure for a sample of trees given a tree metric. We first reduce the set of trees to a pairwise tree distance matrix, with *D*_*i,j*_ = *d*_*RNNI*_ (*t*_*i*_, *t*_*j*_) being the distance between the trees *t*_*i*_ and *t*_*j*_ and *D*_*i*_ denoting the *i*-th row of this matrix. In our experience, considering a single focal tree leads to pseudo ESS estimates that are rather temperamental, sometimes getting stuck at low values depending on the choice of focal tree. Therefore, we consider all trees in the set as focal tree. The pseudo ESS is then defined as the median over the ESS estimate of every row.

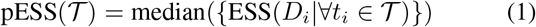

We can use this as a criterium for convergence by making sure pESS(𝒯) is at least as high as the target ESS. Algorithm S1 provides more details.

#### Algorithm S1

Pseudo tree ESS calculation,

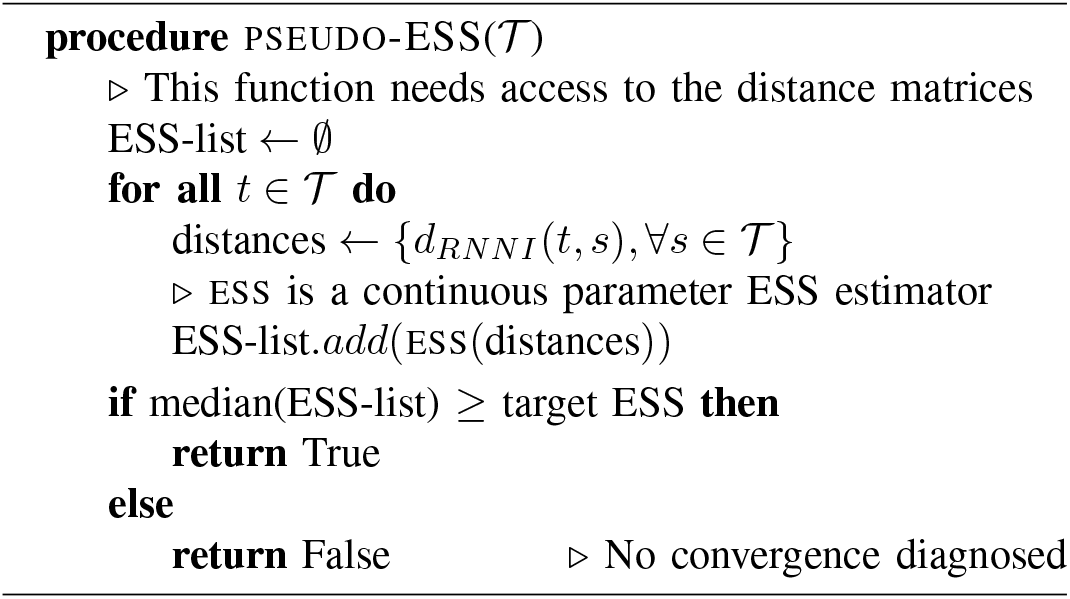

### B Checking ESS of parameter traces

A common convergence check is to make sure ESSs for traces of interest, in particular posterior, prior and likelihood, exceed a certain threshold value. Algorithm S2 shows how this can be automated as a convergence criterion.

#### Algorithm S2

Trace ESS calculation

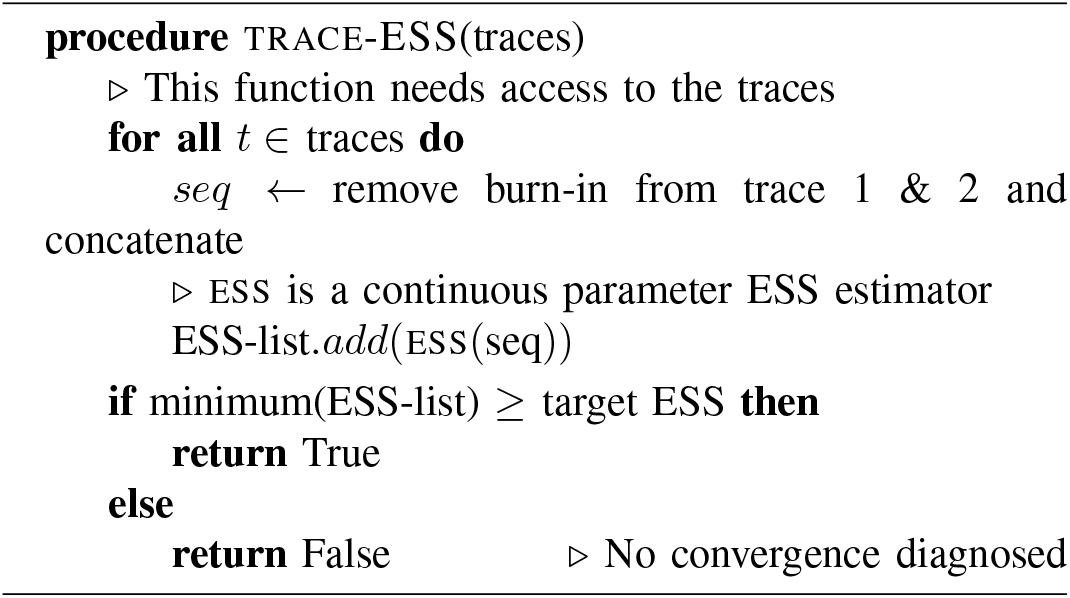

### C Tree PSRF calculation

The following Algorithm S3 presents the calculation of the PSRF value for trees. Its input consists of two sets of trees 𝒯_1_, 𝒯_2_ and an index *t* that is the index of a sample in 𝒯_1_. It returns the PSRF value for the tree in 𝒯_1_ at the given index *t*.

#### Algorithm S3

PSRF calculation for a tree

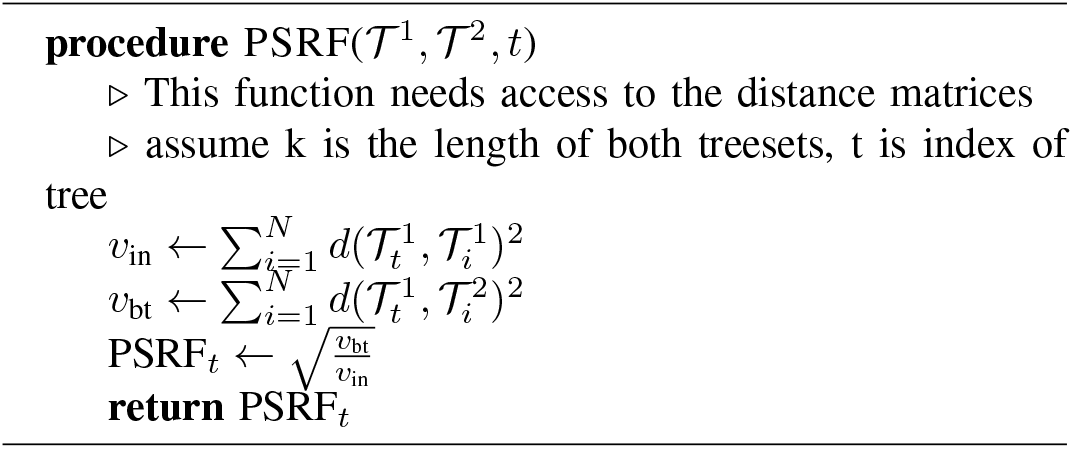

### D GR_T_ diagnostic

This allows us to formulate a stopping criterion presented in Algorithm S4 based on the comprehensive GR_*T*_ diagnostic that we developed. While, for simplicity, we previously excluded the possibility of traces having varying lengths, it is noteworthy that in practice, traces can indeed have different lengths, and burn-in periods between chains can vary significantly. This practical consideration adds to the algorithms’ complexity, leading us to omit it from the pseudo code to enhance readability.

#### Algorithm S4

Automated convergence assessment

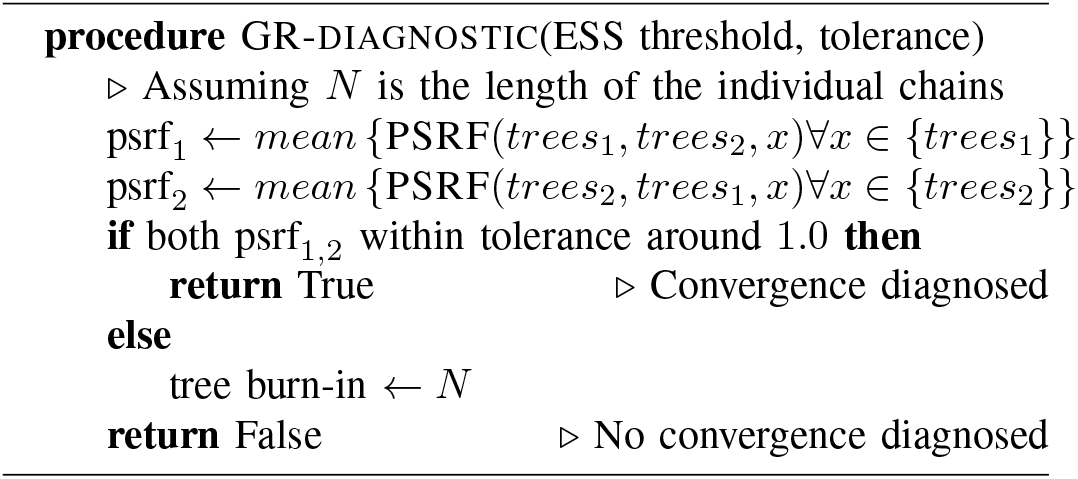

### E Burn-in detection

The following Algorithm S5 estimates the burn-in for a trace of continuous parameters. Its input is a trace *x*_1_, *x*_2_, …, *x*_*n*_ of length *n* and it returns the number of samples that should be discarded as burn-in from the beginning of this trace.

#### Algorithm S5

Burn-in detection

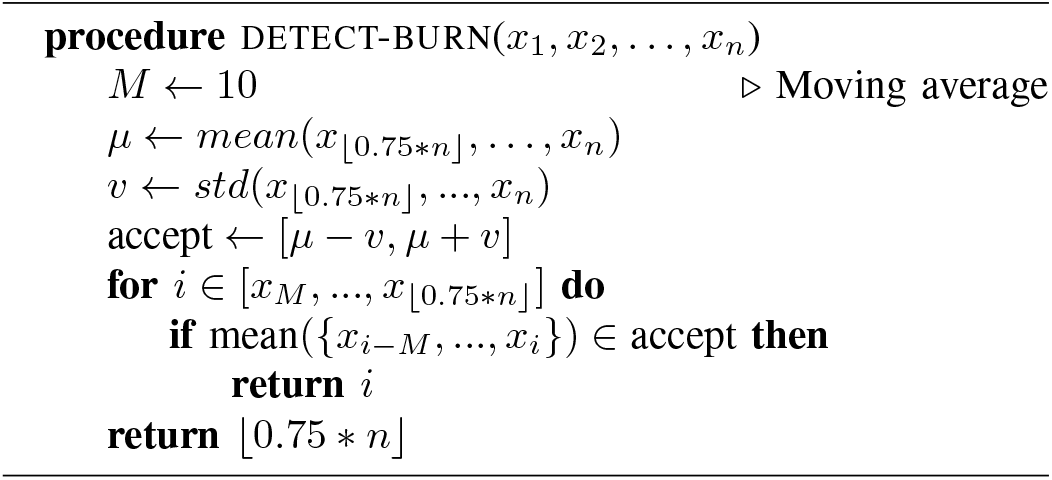

### F Updating matrices

This is an auxiliary algorithm for the GR_*T*_ diagnostic, and is the main additional computational cost at every sampling step of the MCMC. Adding a tree to a distance matrix implies that the distance calculation with complexity *n*^2^, where *n* is the number of taxa, has to be executed *N* times, with *N* being the number of samples of the MCMC that have been stored so far (excluding this newest sample). In addition to adding the trees from each chain to its respective distance matrix we also need to keep track of the in-between distances, which adds additional *N* distance calculations for either tree. Therefore, the added complexity of our diagnostic at every iteration is *O*((4*N* + 1) ∗ *n*^2^).

In practice it is feasible to impose an upper limit on the size of the stored distance matrices. Upon reaching this userdefined limit, every second row and column is removed from the matrix, and not further considered in pseudo-ESS and PSRF calculations (Algorithms S1 and S3). Moreover, future updates exclusively apply to even-numbered trees, with odd numbered trees disregarded. Upon reaching this size limitation *k* times, only trees numbered *i* are considered, where *i* is divisible by 2^*k*^.

#### Algorithm S6

Updating distance matrices

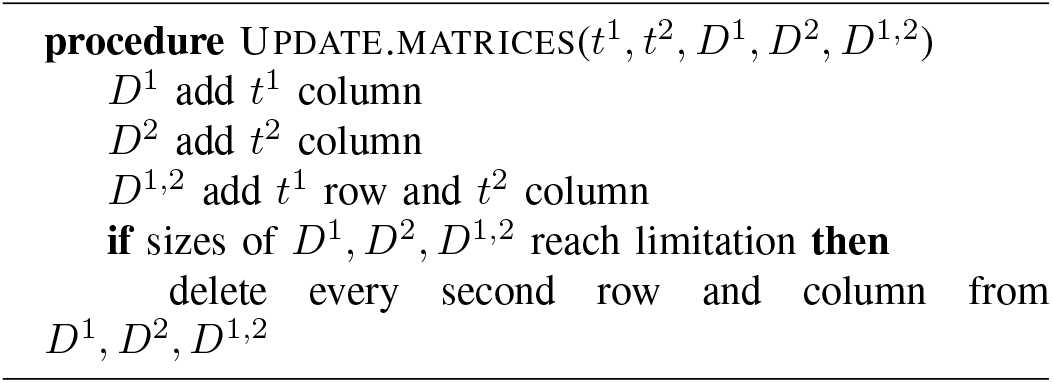

### G Automatically stopping MCMC

The complete automatically stopping MCMC runs two independent MCMC chains, as well as a complete diagnostic tool (Methods section) executed in parallel with the MCMC analyses. The algorithm, detailed in Algorithm S7, is invoked for each iteration of the sampling process to determine the convergence status of the chains. This assessment considers the number of samples, ESS estimates for continuous parameters and trees, and the PSRF values.

#### Algorithm S7

Have the chains converged?

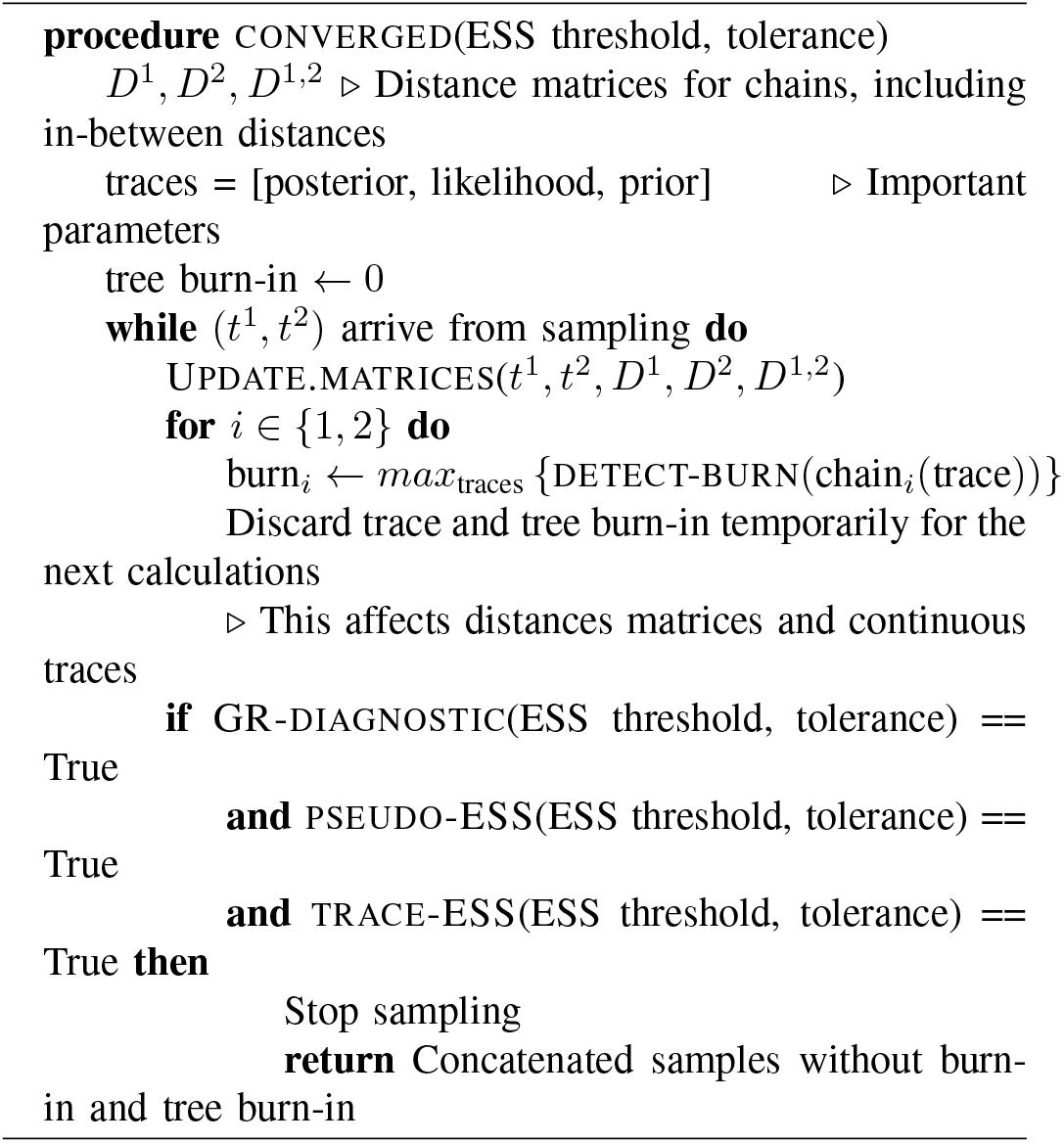

## S. GR VALUE RELATED DECISIONS

In this section we provide insight into why we choose the tolerance of the GR value to be 0.05 or lower. In addition we showcase why the “smoothing” or rather the use of an average function for the GR value is essential.

### A Choice of tolerance

The tolerance of 0.05 is chosen based on the following result, which despite only showing one specific experiment was done for multiple different analyses and datasets. The following plots display the GR value for a fully run analysis, i.e. two independent chains sampled the tree sets 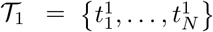 and 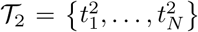 then the values plotted on the y axis are

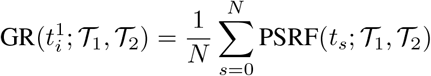

And

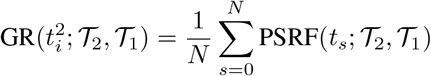

for each sample *i* (x axis).

Such an analysis is visualised in Figure S3 were it is visible that the GR values for almost all samples fall into the interval (0.98, 1.02). The density, displayed in the right plot is very high and showcases how narrow this interval is. In Figure S3, on the other hand, we visualised the same values but using 2 samples of trees from different simulations. It can be observed that these GR values are much higher, have very big variances, and the two samples of trees have very distinct values. This gives us confidence that, due to its sensitivity, our diagnostic is accurately diagnosing convergence problems in treespace. These convergence problems include slowly mixing chains, chains that are stuck or sampling different regions of treespace, or simply chains that are not sampling similar enough trees.

### B Why smoothing is necessary?

Here we present why smoothing of the PSRF values, which results in the GR value, is necessary for our diagnostic. The smoothing takes place whenever the GR value is calculated and we compared using the mean

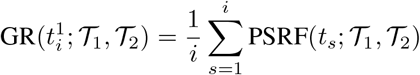

or median

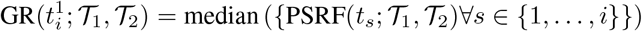

In Figure S4 we display the PSRF values without smoothing, i.e. at every iteration *i* (x axis) the value PSRF(*t*_*i*_; 𝒯_1_, 𝒯_2_) is plotted (y axis). It can be seen that these values have a very high variance and absolute value when compared to the previously presented GR values in Figure S2.

**Fig. S2:**
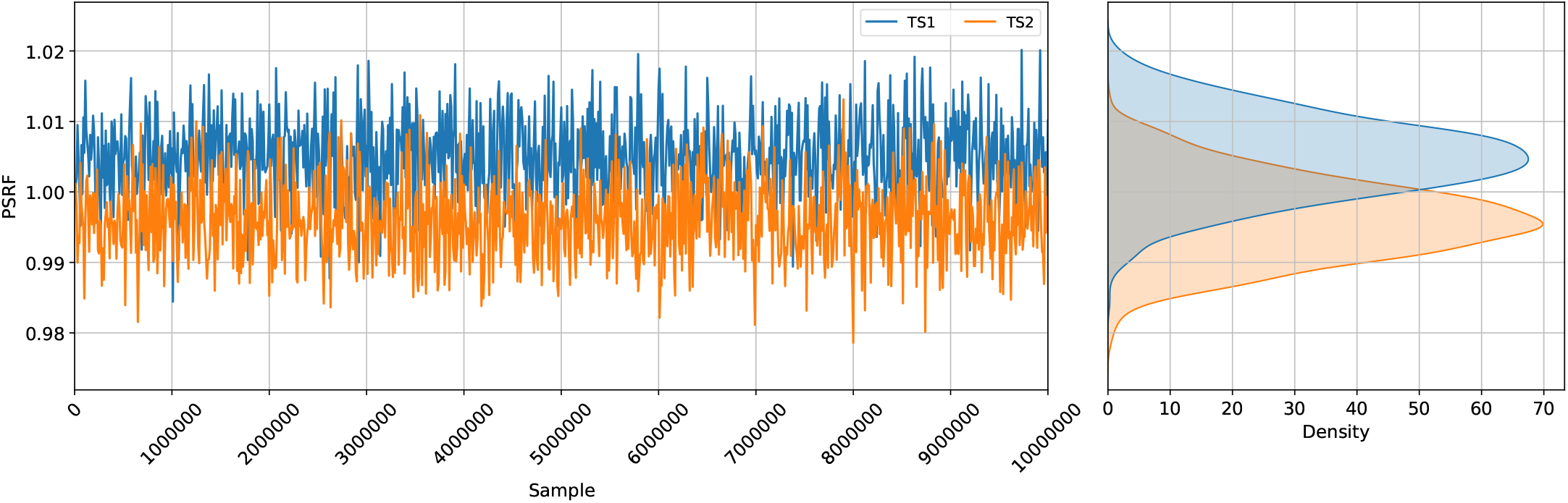
GR value for two independent chains on the same dataset. The left part of the plot displays these values for each sample in the chain and the plot on the right visualises the density of these values for each chain. Notably, the density of these values is very high and the variance around the value 1 is very low.

**Fig. S3:**
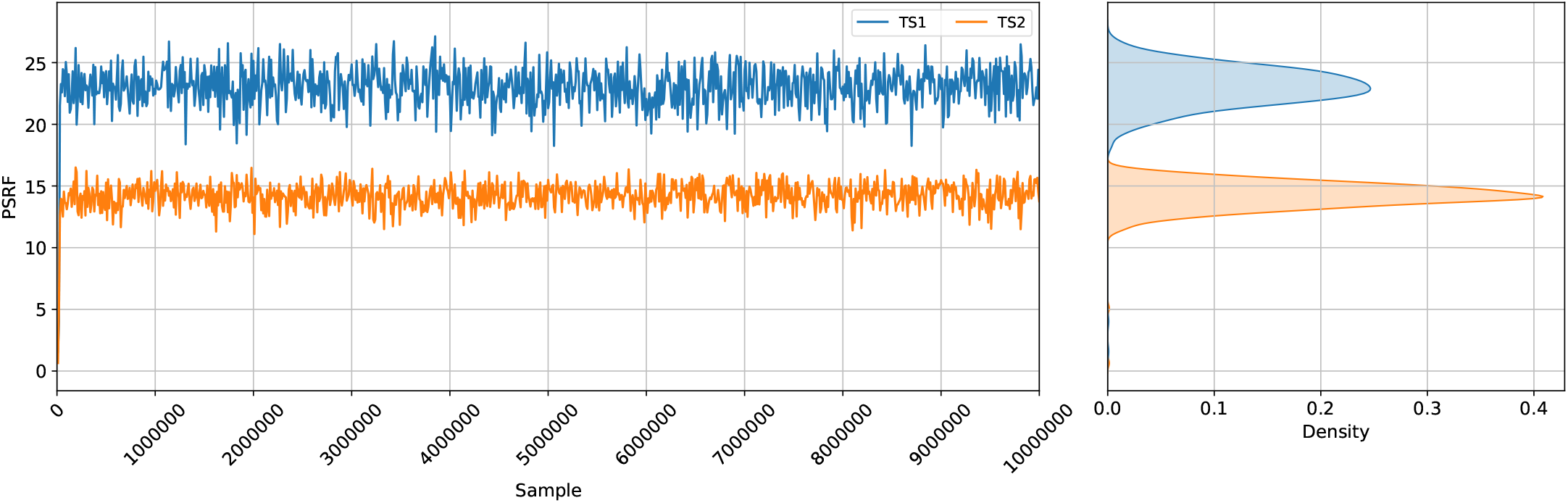
GR value for two independent chains on different datasets. The left displays the GR values for both chains and the right visualises the density for both chains. Unlike the previous plot there is a very wide gap between the two chains and the absolute value is comparably very high, the variance is also very high.

**Fig. S4:**
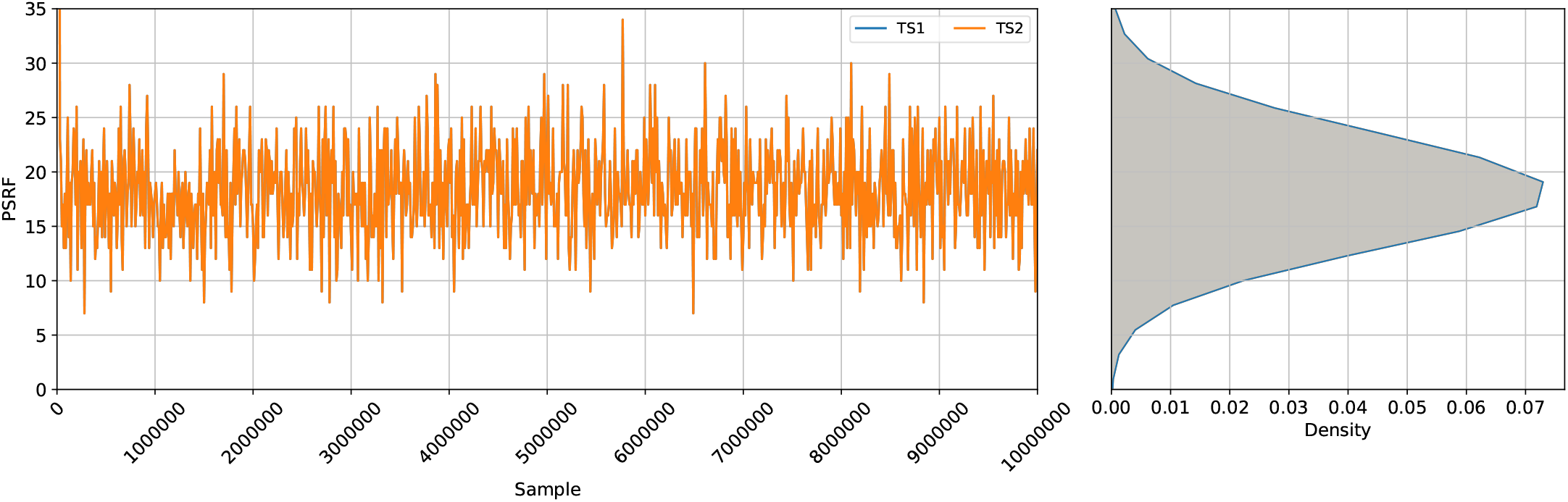
Displaying only the PSRF value at each iteration on the x axis. These have very high absolute values and a large variance (y axis).

## S4 Validation

We compare the use of the median smoothing instead of the mean. Moreover, we provide more results in situations with different GR tolerances and present coverage tables for these different setups of our diagnostic.

### A Comparing mean and median convergence

In Table S1 we highlight the difference when using the median instead of mean to smooth out the PSRF values. There are no significantly big differences among the two averages, the median could be seen as a slightly more conservative measurement.

**TABLE S1:** Our convergence diagnostic as implemented in tetres executed on 100 well-calibrated simulation studies with 1000 trees each. Highlighted in red is the only difference (2 instead of 1) between using the median and mean to average the PSRF values for the calculation of the GR value.

| Tolerance | ESS threshold | Converged | Not converged |
| --- | --- | --- | --- |
| 0.05 | 200 | 100 | 0 |
|  | 500 | 100 | 0 |
| 0.02 | 200 | 98 | 2 |
|  | 500 | 92 | 8 |
| 0.01 | 200 | 80 | 20 |
|  | 500 | 53 | 47 |

#### a Coverage analyses

The following tables include coverage analysis when using different tolerances. Note, that not all of the 100 runs are considered converged under different settings, therefore each column has individual number of total simulations and corresponding confidence intervals.

**TABLE S2:** mean-0.02.

| Paramter | ESS threshold, total runs, confidence interval |  |  |
| --- | --- | --- | --- |
|  | 200, 99, (90-98) | 500, 92, (84-91) | Full, 100, (91-99) |
| Tree.height | 92/91 | 88/88 | 96/96 |
| Tree.treeLength | 93/92 | 87/86 | 97/94 |
| clockRate | 89/91 | 83/85 | 91/93 |
| gammaShape | 88/89 | 85/84 | 92/92 |
| popSize | 93/91 | 89/89 | 96/97 |
| CoalescentConstant | 91/89 | 85/84 | 92/92 |
| kappa | 90/95 | 85/86 | 94/94 |
| freqParameter.1 | 91/91 | 85/85 | 92/92 |
| freqParameter.2 | 89/91 | 85/87 | 94/95 |
| freqParameter.3 | 86/85 | 81/76 | 87/90 |
| freqParameter.4 | 95/93 | 84/84 | 95/95 |

**TABLE S3:** mean-0.01.

| Paramter | ESS threshold, total runs, confidence interval |  |  |
| --- | --- | --- | --- |
|  | 200, 80, (72-79) | 500, 53, (48-53) | Full, 100, (91-99) |
| Tree.height | 74/76 | 51/50 | 96/96 |
| Tree.treeLength | 76/75 | 49/50 | 97/94 |
| clockRate | 73/72 | 49/49 | 91/93 |
| gammaShape | 72/71 | 49/49 | 92/92 |
| popSize | 76/75 | 51/50 | 96/97 |
| CoalescentConstant | 73/73 | 48/48 | 92/92 |
| kappa | 73/75 | 49/49 | 94/94 |
| freqParameter.1 | 72/74 | 49/49 | 92/92 |
| freqParameter.2 | 73/76 | 48/49 | 94/95 |
| freqParameter.3 | 69/70 | 45/45 | 87/90 |
| freqParameter.4 | 73/74 | 48/49 | 95/95 |

### B Median smoothing coverage for well-calibrated simulation study

The following tables display the coverage analyses when using the median smoothing option for the GR value. Because of no notable difference to the version using the mean, the coverage tables are also very similar. Note, that not all of the 100 runs are considered converged under different settings, therefore each column has individual number of total simulations and corresponding confidence intervals.

**TABLE S4:** median-0.05.

| Paramter | ESS threshold, total runs, confidence interval |  |  |
| --- | --- | --- | --- |
|  | 200, 100, (91-99) | 500, 100, (91-99) | Full, 100, (91-99) |
| Tree.height | 92/94 | 95/96 | 96/96 |
| Tree.treeLength | 94/92 | 96/95 | 97/94 |
| clockRate | 91/92 | 90/92 | 91/93 |
| gammaShape | 87/90 | 92/93 | 92/92 |
| popSize | 90/94 | 97/95 | 96/97 |
| CoalescentConstant | 91/91 | 93/91 | 92/92 |
| kappa | 92/94 | 93/93 | 94/94 |
| freqParameter.1 | 94/91 | 92/92 | 92/92 |
| freqParameter.2 | 93/92 | 94/94 | 94/95 |
| freqParameter.3 | 86/87 | 89/88 | 87/90 |
| freqParameter.4 | 95/95 | 94/94 | 95/95 |

**TABLE S5:** median-0.02.

| Paramter | ESS threshold, total runs, confidence interval |  |  |
| --- | --- | --- | --- |
|  | 200, 98, (89-97) | 500, 92, (84-91) | Full, 100, (91-99) |
| Tree.height | 91/90 | 88/88 | 96/96 |
| Tree.treeLength | 91/92 | 88/86 | 97/94 |
| clockRate | 89/89 | 83/84 | 91/93 |
| gammaShape | 86/89 | 85/82 | 92/92 |
| popSize | 92/93 | 89/89 | 96/97 |
| CoalescentConstant | 90/88 | 85/84 | 92/92 |
| kappa | 89/92 | 85/86 | 94/94 |
| freqParameter.1 | 90/90 | 85/85 | 92/92 |
| freqParameter.2 | 89/90 | 85/87 | 94/95 |
| freqParameter.3 | 85/83 | 81/77 | 87/90 |
| freqParameter.4 | 93/92 | 85/84 | 95/95 |

**TABLE S6:** median-0.01.

| Paramter | ESS threshold, total runs, confidence interval |  |  |
| --- | --- | --- | --- |
|  | 200, 80, (72-79) | 500, 53, (48-53) | Full, 100, (91-99) |
| Tree.height | 75/76 | 51/50 | 96/96 |
| Tree.treeLength | 75/75 | 49/50 | 97/94 |
| clockRate | 73/73 | 49/49 | 91/93 |
| gammaShape | 73/72 | 49/49 | 92/92 |
| popSize | 77/77 | 51/50 | 96/97 |
| CoalescentConstant | 72/72 | 48/48 | 92/92 |
| kappa | 73/74 | 49/49 | 94/94 |
| freqParameter.1 | 73/75 | 49/49 | 92/92 |
| freqParameter.2 | 73/75 | 48/49 | 94/95 |
| freqParameter.3 | 69/70 | 45/45 | 87/90 |
| freqParameter.4 | 72/73 | 48/50 | 95/95 |

## S5. Beast ASM package on DS1–DS11 and other data sets

The following Table S7 displays the individual run lengths of 10 independent chains on HCV, primate and COVID-19 alignments using the ASM package, each chain has a sample frequency of 10, 000.

**TABLE S7:**
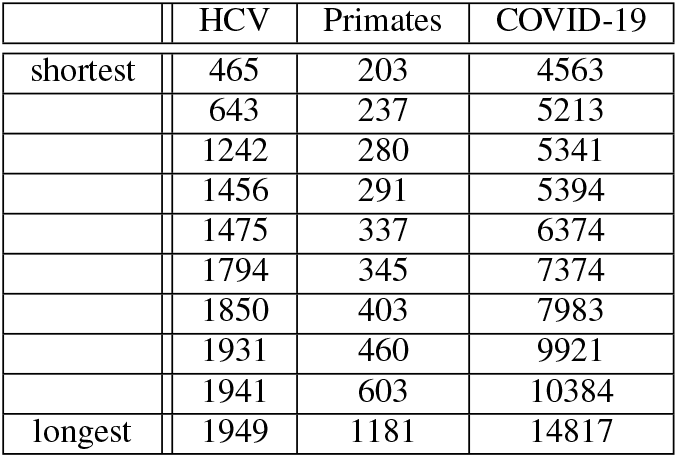
MCMC chain lengths (divided by 10^4^) over 10 runs before automatically stopping j.

|  | HCV | Primates | COVID-19 |
| --- | --- | --- | --- |
| shortest | 465 | 203 | 4563 |
|  | 643 | 237 | 5213 |
|  | 1242 | 280 | 5341 |
|  | 1456 | 291 | 5394 |
|  | 1475 | 337 | 6374 |
|  | 1794 | 345 | 7374 |
|  | 1850 | 403 | 7983 |
|  | 1931 | 460 | 9921 |
|  | 1941 | 603 | 10384 |
| longest | 1949 | 1181 | 14817 |

The following tables display the individual run lengths of 10 independent chains on DS1–DS11 using the ASM package, each chain has a sample frequency of 10, 000.

**TABLE S8:**
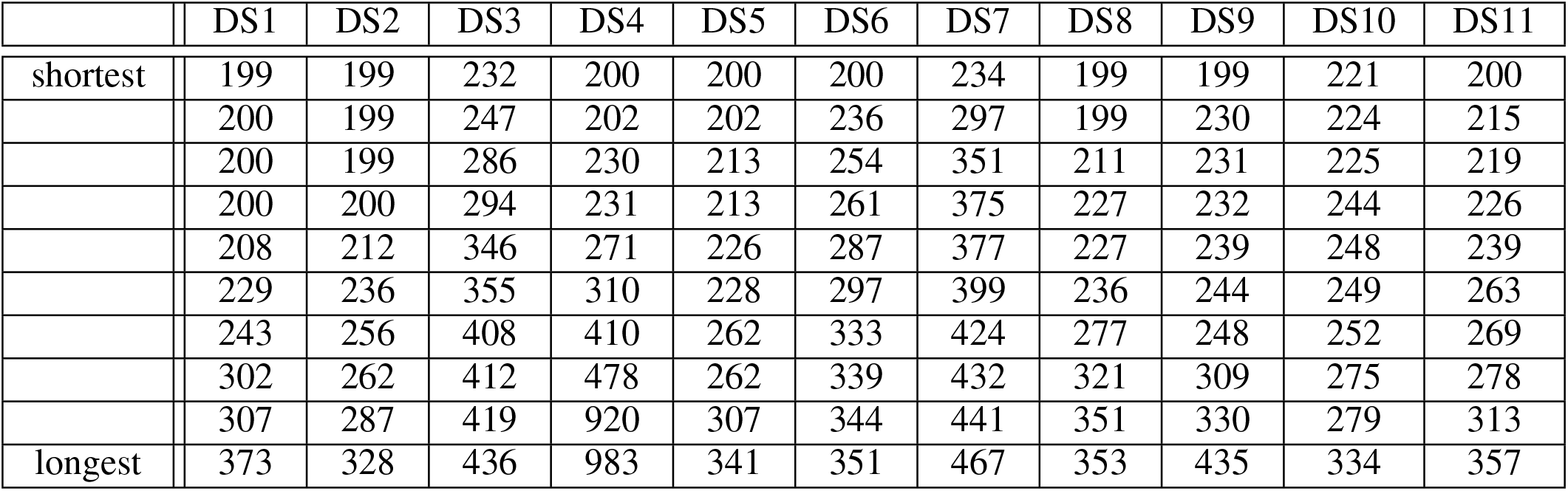
MCMC chain lengths (divided by 10^4^) over 10 replicates before automatically stopping.

|  | DS1 | DS2 | DS3 | DS4 | DS5 | DS6 | DS7 | DS8 | DS9 | DS10 | DS11 |
| --- | --- | --- | --- | --- | --- | --- | --- | --- | --- | --- | --- |
| shortest | 199 | 199 | 232 | 200 | 200 | 200 | 234 | 199 | 199 | 221 | 200 |
|  | 200 | 199 | 247 | 202 | 202 | 236 | 297 | 199 | 230 | 224 | 215 |
|  | 200 | 199 | 286 | 230 | 213 | 254 | 351 | 211 | 231 | 225 | 219 |
|  | 200 | 200 | 294 | 231 | 213 | 261 | 375 | 227 | 232 | 244 | 226 |
|  | 208 | 212 | 346 | 271 | 226 | 287 | 377 | 227 | 239 | 248 | 239 |
|  | 229 | 236 | 355 | 310 | 228 | 297 | 399 | 236 | 244 | 249 | 263 |
|  | 243 | 256 | 408 | 410 | 262 | 333 | 424 | 277 | 248 | 252 | 269 |
|  | 302 | 262 | 412 | 478 | 262 | 339 | 432 | 321 | 309 | 275 | 278 |
|  | 307 | 287 | 419 | 920 | 307 | 344 | 441 | 351 | 330 | 279 | 313 |
| longest | 373 | 328 | 436 | 983 | 341 | 351 | 467 | 353 | 435 | 334 | 357 |

